# Antimicrobial peptides expressed by plant-engineered ‘symbiont’ technology reduces titers and disease symptoms of “*Candidatus* Liberibacter solanacearum” in potato

**DOI:** 10.64898/2026.08.07.743526

**Authors:** W. Rodney Cooper, Laura A. Fleites, Robert G. Shatters, Marco Pitino, Samuel Coradetti, Michelle Heck

## Abstract

Delivery of therapeutic biomolecules into plant vascular tissues remains a challenge in management of vector-borne plant pathogens. The symbiont concept uses reprogrammed *Agrobacterium tumefaciens* galls (called symbionts) to produce biomolecules while remaining connected to host vasculature. We evaluated whether symbionts expressing antimicrobial peptides (AMPs) suppress ‘*Candidatus* Liberibacter solanacearum’ (CLso), the causal agent of potato zebra chip disease. Symbionts were engineered to express a *Streptococcus mutans* bacteriocin associated with bacterial membrane disruption (Blp-Sm), or an AMP isolated from finger lime and associated with resistance to citrus greening disease (MaSAMP). Effects of AMP-producing symbionts on CLso titers, infection incidence, pathogen movement, and disease symptoms were evaluated in tomato and potato. In tomato, neither AMP significantly reduced CLso titers or infection incidence. However, in potato, AMP-producing symbionts reduced CLso accumulation and movement from CLso-inoculated source shoots into non-inoculated sink shoots connected through underground tubers. Blp-Sm produced the strongest reduction in CLso accumulation and infection incidence in sink tissues. In separate assays where symbionts were established directly on potato seed tubers, MaSAMP significantly reduced CLso titers in stems and tubers and reduced zebra chip symptoms in tubers, despite no reduction of CLso titers in terminal leaves. These findings demonstrate that AMP-producing symbionts suppress vascular pathogen accumulation and movement within plants and highlight the symbiont concept as a potential platform for managing diseases caused by vascular-restricted pathogens. Further, they show the potato-CLso system is a promising infection model to both refine and improve symbiont technology, and to test additional AMPs for potency against related pathogens.

## Introduction

Delivery of therapeutic biomolecules into plant vascular systems remains a major obstacle in managing insect-vectored plant pathogens such as *Candidatus* Liberibacter spp. and *Ca.* Phytoplasma spp., which cause severe economic losses and threaten entire agricultural commodities. For example, *Ca.* Liberibacter asiaticus (CLas), the presumed causal agent of citrus greening disease (Huanglongbing), has contributed to a 95% decline in Florida citrus production from 2004 to 2025 and more than $75 billion in economic damage to the U.S. citrus industry. Similarly, *Ca.* Phytoplasma pruni, responsible for cherry X-disease, has caused approximately $30 million in annual losses to the Pacific Northwest cherry industry since 2015 (Harper et al. 2023). Effective management is hindered by the fact that these pathogens often remain asymptomatic for 2–4 years in perennial crops while highly mobile insect vectors continue to spread infections to healthy trees and orchards (Lee et al. 2015). Growers rely heavily on repeated insecticide applications to suppress vector populations, yet these measures have little to no impact on preventing pathogen spread. With no curative treatments available, infected trees must be removed once symptoms appear. There is a critical need for alternative tools that can directly target and control vascular-restricted plant pathogens.

A novel approach to plant bioengineering with the potential to deliver biomolecules directly to pathogens residing in plant phloem or xylem tissue involves the use of transgenic, non-pathogenic galls, referred to as symbionts, that express therapeutic peptides (Heck et al. 2026). The symbiont approach relies on expression of the *Agrobacterium tumefaciens* plant growth regulator gene cassette together with a gene of interest, such as those that encode for antimicrobial peptides, together on the same piece of transfer DNA (T-DNA). Thus, symbionts are reprogrammed host plant cells that form a semi-organized external gall-like structure. The goal of the symbiont concept is to deliver biomolecules into the plant’s vascular tissues capable of modifying whole-plant phenotypes in established trees or crops (Heck et al. 2026); while the rest of the tree is not genetically engineered. The symbiont concept has been demonstrated on herbaceous and woody dicots by expression of fluorescent reporters or RNA-silencing constructs, which accumulated in and exported from the symbiont galls into adjacent tissues (Heck et al. 2026). Formation and growth of symbiont galls on plant stems did not significantly negatively impact plant growth and fruit yield. Although expression of marker proteins by symbiont cell clusters provides a proof-of-concept for symbiont biotechnology, delivery of antimicrobial biomolecules for pathogen suppression has not yet been demonstrated.

The goal of our study was to evaluate the impact of symbiont technology on pathogen performance by delivery of antimicrobial peptides (AMPs). We used *Ca.* Liberibacter solanacearum (CLso) and its vector, *Bactericera cockerelli* (potato psyllid) as a study system. CLso infects plants within the Solanaceae, causing zebra chip disease of potato and foliar disease symptoms in other Solanaceous crops including tomato and pepper (Wenninger and Rashed 2024). CLso titers and disease symptoms develop rapidly in potato and tomato and are measurable several weeks after initial infection (Sengoda et al. 2010, 2014; Cooper et al. 2014).

We evaluated the impact of two different AMPs expressed in symbiont that were previously identified as potential CLas active candidates in a preliminary screening program (unpublished results from Dr. Shatters’ laboratory): 1) a bacteriocin identified from *Streptococcus mutans* (Dawid et al. 2007, Franz et al. 2007) hereafter referred to as Blp-Sm, or 2) a stable antimicrobial peptide identified from finger lime, *Microcitrus australiasica*, hereafter referred to as MaSAMP (Haung et al. 2021). Bacteriocins are antimicrobial peptides produced by bacteria to support niche competition with related bacteria and are generally grouped into three main classes (and a fourth less common) for gram positive bacteria: <u>Class I lantibiotics</u>, which are small, heat-stable peptides containing unusual amino acids like lanthionine; <u>Class II non-lantibiotics</u>, also small and heat-stable but lacking unusual residues and further divided into subgroups such as pediocin-like, two-peptide, cyclic, and miscellaneous bacteriocins; and <u>Class III bacteriocins</u>, which are large, heat-labile proteins that often function as bacteriolytic enzymes.

Some systems also include <u>Class IV</u>, complex bacteriocins requiring lipid or carbohydrate components. <u>Gram-negative systems</u> also produce bacteriocins including colicin-and microcin-types, but the first three classes remain the most commonly studied. Blp-Sm belongs to the class II group of bacteriocins that exhibit broad-spectrum activity against bacteria by forming pores in the bacterial cell membrane leading the cell death (Dawid et al. 2007, Kumariya et al. 2019).

The Stable Antimicrobial Peptide (SAMP) from Australian finger lime (*Microcitrus australasica*) is a small (∼6.7 kDa) heat-stable AMP that exhibits dual functionality (Huang et al. 2021 and Wang et al. 2021): it both directly kills CLas and activates plant innate immunity. Since it is from *M. australasica* we will refer to it as Ma-SAMP). The short (∼6. kDa) isoform accumulates in the phloem, where CLas resides, and its α-helix-2 domain is sufficient to kill Liberibacter, causing rapid cytosolic leakage and cell lysis. Greenhouse trials showed that foliar or trunk application significantly reduces CLas titers and HLB symptoms in infected trees, while also boosting resistance to new infection in healthy trees. We demonstrate that both Blp-Sm and MaSAMP expressed in symbiont reduce CLso titers in tissues adjacent to transformed tissues and inhibits the vascular transport of CLso through stems with symbiont galls structures.

## Materials and Methods

### Sources of plants and CLso-infected *B. cockerelli*

Potato (cultivar “Ranger Russet”) and tomato (cultivar “Micro Tom”) plants were grown in a greenhouse at ∼25°C with supplemental lighting to provide a 16:8 h photoperiod. Potatoes and tomatoes were grown in 1.5-l and 0.5-l pots, respectively, filled with Miracle Grow potting mix (Scotts Miracle Gro Inc., Marysville, OH). *Bactericera cockerelli* and CLso occur as biologically distinct haplotypes (Swisher et al. 2012). Psyllids of the western haplotype maintained in a laboratory colony on tomato and potato were used for all assays because they readily occur in potato fields and because they efficiently transmit CLso to uninfected plants (Swisher et al. 2014; Cooper et al. 2015). CLso haplotype-B was used for all assays because it is highly virulent to plants (Mendoza-Herrera et al. 2018; Swisher Grimm et al. 2018; Reyes Corral et al. 2020). The colonies were tested monthly using standard diagnostic PCR (Crosslin et al. 2011) to confirm CLso infection.

### Symbiont construction and inoculation

The pSym plasmid was constructed as described in Heck et al. (2026). Its sequence is available on GenBank (PQ563460) and can be obtained from the global, non-profit microbial resource center, ATCC (Deposition # MBA-395). Briefly, the PGR cassette from *A. tumefaciens* strain C58 was PCR-amplified and ligated into pUSHRL026 at the Ascl site immediately upstream of the double 35S promoter. Symbiont constructs were built to express the Blp-Sm or MaSAMP with GFP as a marker protein. A vector expressing the antimicrobial peptide, pSym-GFP-AMP, was constructed by fusing the AMP coding sequence to the C-terminus of GFP under the 35S promoter, with a P2A ribosomal skipping site (Wang and Marchisio 2021). The placement of these AMPs immediately after a P2A site resulted in a single proline addition to the N-terminus of those AMPs. EHA105 carrying pSym plasmids were grown overnight in LB with 50 μg/ml kanamycin at 28°C at 180 rpm until the culture. The liquid cultures were centrifuged to pellet the cells, which were washed three times with 10 ml of inoculation solution (10mM MgCl2, 10mM MES, and 500μM acetocyringone). The cells were resuspended in 10 ml of inoculation solution and adjusted to 1.0 OD_600_.

### AMPs expressed from stem-inoculated symbiont on tomato and potato

Two assays were developed to test the effects of symbiont-delivered AMPs on CLso on potato and tomato stems (**Fig. 1A**). The tomato stem assay tested the impact of symbionts directly on the CLso inoculated shoot. The potato assay tested for symbiont effects on CLso movement between two shoots arising from the same mother tuber (**Fig. 2**).

**Figure 1.**
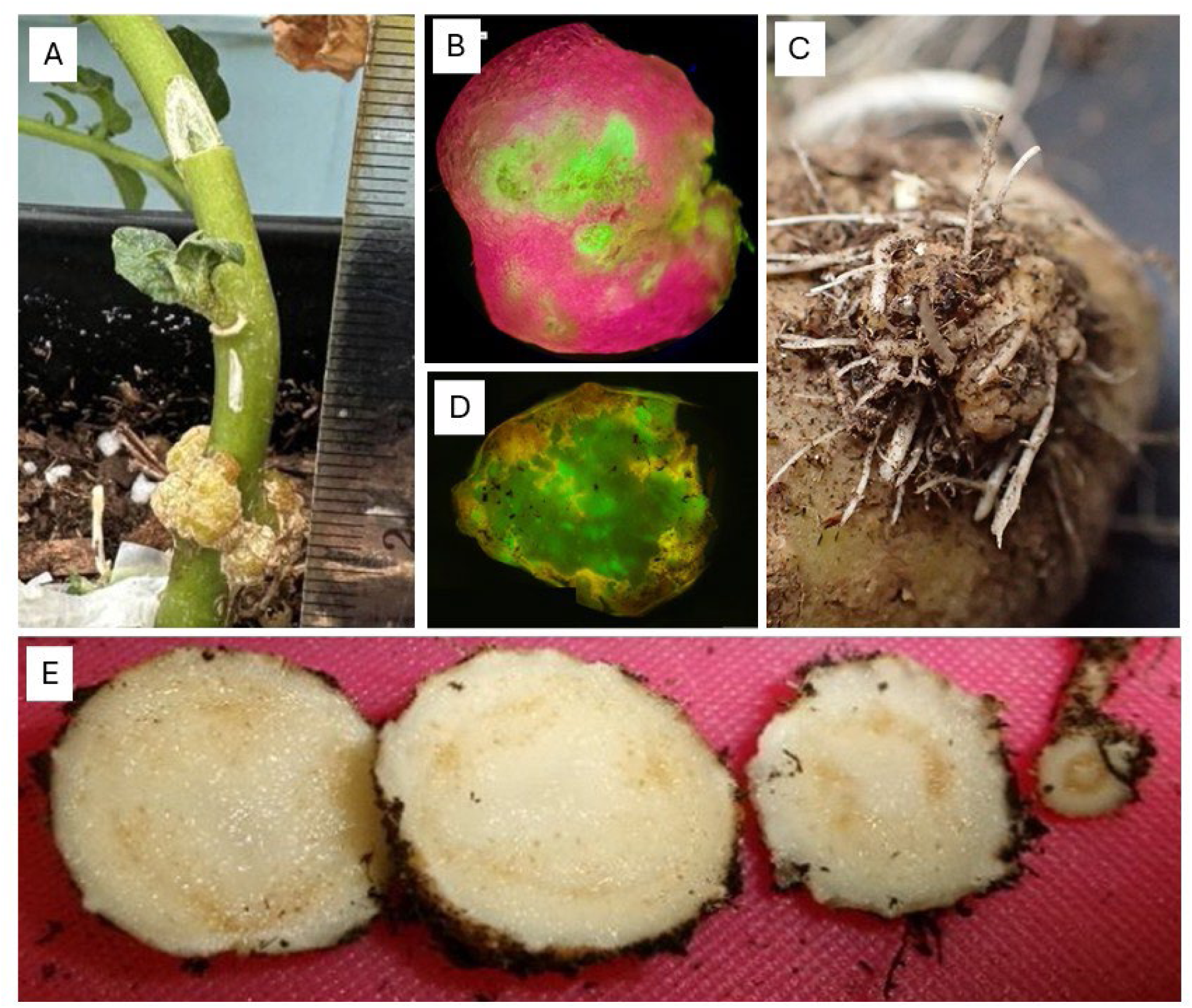
Symbiont gall growing on a potato stem with tissue extracted above the symbiont for CLso quantification (A). GFP expression in a symbiont grown on a tomato stem (B). Symbiont gall growing on a potato seed tuber (C) and observed GFP expression in the sliced symbiont (D). Zebra chip symptoms observed in a sliced daughter tuber (E).

**Figure 2.**
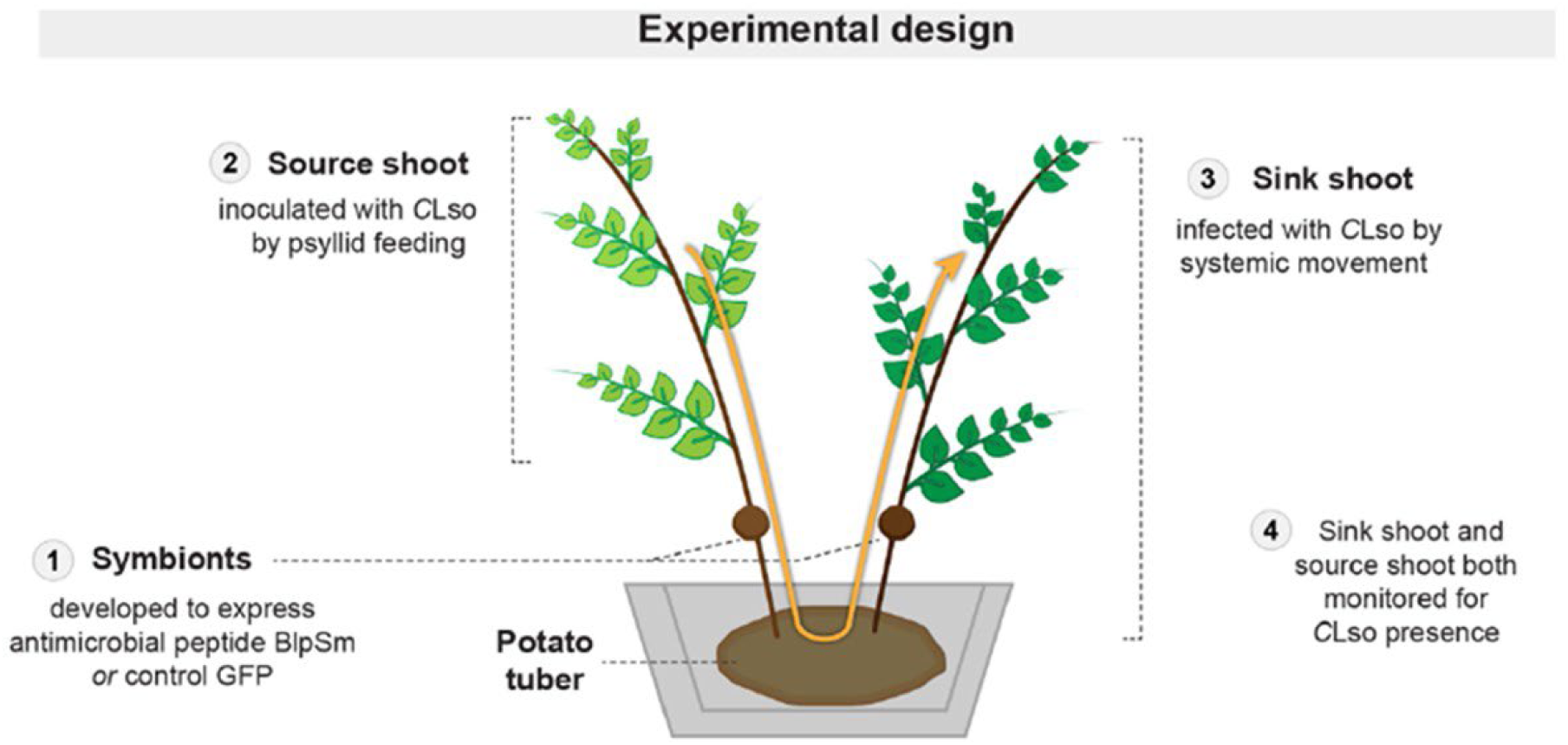
Experimental design for potato stem assay.

In the tomato stem assay, the impact of AMP-expressing symbionts on CLso infection rates and titers was first examined on tomato stems. Blp-Sm and MaSAMP were tested in separate independent experiments by comparing CLso titers and infection rates in plants with AMP-producing symbionts versus symbionts expressing GFP only. Plant stems were inoculated with symbiont-forming *A. tumefaciens* using a 27G insulin syringe 2-3 cm above the soil line. The needle was pushed through the entire stem while solution was gently injected as the needle was removed. The inoculation was repeated by injecting the stem at a perpendicular angle directly above the initial inoculation site. This process produced four symbiont growths on each stem (**Fig. 1A**). After inoculation, the plants were placed in a growth room maintained at 22°C with a 16-h photoperiod and 60% humidity for cell transformation. Four weeks after inoculation, each plant was infected with CLso by confining six colony-reared psyllids to the 3^rd^ leaf from the soil using sleeve cages. Psyllids were removed from plants after 1 week by excising the caged leaf, and CLso infection was confirmed from each group of psyllids using conventional PCR described below. Plants were moved either to a greenhouse or a growth chamber with 40% humidity to prevent stomatal dilation and bursting. The terminal leaves and stem tissue (3 cm above the gall; **Fig. 1A**) were collected from plants and stored in RNA*later* (Invitrogen, Waltham, MA) in - 80°C until processed for CLso quantification. Each gall was sliced and examined for GFP expression (**Fig. 1B**) using a handheld NightSea Fluorescence Flashlight Xite-RB with an excitation wavelength of 440-460 nm and a barrier filter glass of 500 nm (NightSea, Hatfield, PA).

The potato stem assay tested for symbiont effects on the systemic movement of CLso in plants. Potato plants produce multiple shoots that arise from a common seed tuber (mother tuber) beneath the soil (**Fig. 2**). CLso readily moves between stems though the mother tuber (Levy et al. 2011), providing a means to examine how AMP-expressing symbionts influence movement of CLso in plants. The impact of Blp-Sm and MaSAMP on CLso infecting potato was examined in separate experiments. Each potato plant was pruned to include just two 10-15 cm shoots, and each shoot was inoculated with symbiont-forming *A. tumefaciens* (**Fig. 2**). One shoot (source shoot) from each plant was infected with CLso four weeks after symbiont inoculation, while the remaining shoot (sink) was used to examine movement of CLso within plants (**Fig. 2**). In this manuscript, our use of the terms source and sink refer to the CLso-inoculated (source) and non-inoculated (sink) stems from a single mother tuber, as illustrated schematically in the experimental design workflow (Fig. 2). Stem and leaf tissues were collected to measure CLso titers and incidence and symbionts were examined for GFP expression as described for tomato (**Fig. 1B**).

All stem-inoculated symbiont experiments were conducted in at least two independent trials as follows. Blp-Sm was tested on tomato in three trials each with 10-15 reps (plants) per treatment and on potato in three trials each with 10 reps per treatment. MaSAMP was tested on tomato in two trials each with 15 reps per treatment and on potato in two trials each with 10 reps per treatment. All data were analyzed using the GLIMMIX procedure of SAS 9.4 (SAS Institute, Cary, NC). Analyses of assays on tomato included treatment as the independent variable. Analyses of experiments using potato included treatment, shoot, and the main effect interaction as independent variables. Infection rates of leaves and stems were analyzed separately using logistic regression by including the LINK=LOGIT, DDFM=NONE, and CHISQ options of the MODEL statement. CLso titers (copies/1000) were analyzed assuming a gamma distribution using the DIST=GAMMA option of the model statement and therefore excluded values of zero titers. Tukey adjustments were used to compare means when analyses indicated a significant main effect interaction.

### AMPs expressed from symbionts located on underground potato seed tubers

A potato tuber assay was also developed to test for symbiont effects when developed directly on seed tubers. Seed tubers of potato Ranger Russet were held for 48 hours at room temperature to break tuber dormancy and initiate tuber bud (eye) swelling. Three eyes on each tuber were inoculated with symbiont-producing *A. tumefaciens* using a 27-G needle. Symbiont treatments included Blp-Sm, MaSAMP, or GFP only (control). The symbiont-inoculated tubers were held in dark within a growth chamber maintained at 25°C with 60% relative humidity for 1 week before planting in soil. When plants reached a 10-15 cm height, they were infected with CLso by confining 6 infected psyllids to the third leaf from the soil for 1 week using a sleeve cage. Terminal leaves and stems above the soil line were collected from each infected shoot for CLso quantification after 4 weeks. In addition, tubers were removed from the soil, and three tuber buds (eyes) were collected from each mother tuber for CLso quantification. Galls were examined for GFP expression as described above (**Fig. 1C-D**). In addition, each seed and daughter tuber was sliced at the stem end to record the presence or absence of zebra chip symptoms (**Fig. 1E**). The experiment was conducted twice; trial 1 included five replicate plants per treatment, and trial 2 included 10 replicate plants per treatment.

Data were analyzed using the GLIMMIX procedure of SAS 9.4. CLso titers (copies/1000) in leaf, stem, and eye tissues were analyzed separately with treatment as the independent variable assuming a gamma distribution (DIST=GAMMA option of the MODEL statement). Infection rates were analyzed using logistic regression by including a binary response. Tuber symptoms were analyzed by logistic regression using an events/trials syntax where events=number of symptomatic tubers and trials=number of tubers from each plant. The LINK=LOGIT, CHISQ, and DDFM=NONE options of the MODEL statement was included in each logistic regression analysis.

### Diagnostic PCR for CLso

DNA was extracted from each group of three psyllids using a modified CTAB method (Crosslin et al. 2011). The primers OA2 (GCG CTT ATT TTT AAT AGG AGC GGC A) and OI2c (GCC TCG CGA CTT CGC AAC CCA T) were used to detect the presence or absence of CLso using conventional PCR (Jagoueix et al. 1996). PCR conditions included 35 cycles of 94°C for 30 s, 66°C for 30 s, and 72°C for 60 s. Visible amplicons were observed on 1% agarose gels stained with GelRed Nucleic Acid Stain (Sigma-Aldrich, Inc. St. Louis, MO).

RNA is more abundant than DNA and degrades faster than DNA after cell death. We therefore measured RNA expression as a proxy for pathogen titers because the assay is more sensitive and more directly detects living bacteria cells. A maximum of 100 mg of plant tissue from each sample was chilled in liquid nitrogen and ground to a fine powder using a Tissue Lyser II (Qiagen, Venlo, Netherlands). RNA was extracted from samples using a Qiagen Rneasy Plant Kit following the manufacturer’s protocol and was converted to cDNA using a Qiagen QuantiTect Reverse Transcription Kit. Concentrations were standardized to 20ng/μL. Titers of CLso were measured using qPCR using primers LsoF (CGA GCG CTT ATT TTT AAT AGG AGC) and HLBr (GCG TTA TCC CGT AGA AAA AGG TAG) (Li et al. 2009). Each reaction consisted of 10 μL of 2X SYBR Green Master Mix (Roche), 8 μl PCR grade water, 0.5 μl of each primer (final concentration of 125 nM), and 1 μL of template cDNA. PCR conditions included 10 min at 95°C, 45 cycles of 95°C for 10 sec, 60°C for 10 sec, 72°C for 30 sec, and a final melt curve. The melt curves were analyzed at the end of each run to ensure there was no off-target amplification. Cq values were compared to those of serial dilutions of plasmid standards to estimate the number of copies of Lso present in each sample.

## Results

### Symbiont production of antimicrobial peptides did not significantly reduce CLso infection rates in tomato

Symbiont gall sizes on tomato plants on average grew to a diameter of 6 mm and consistently expressed GFP (**Table 1**). The presence of symbionts constructed to produce Blp-Sm did not reduce CLso titers in plant stems (**Fig. 3A**; *F*=0.9; df=1, 41; *P*=0.335) or CLso infection rates (**Fig. 3B**; χ^2^=1.4; d.f.=1; *P*=0.236) on tomato. Similarly, the presence of symbionts constructed to produce MaSAMP did not reduce titers in plants stems (**Fig. 4A**; *F*=0.6; df=1, 27; *P*=0.804). Although infection rates were more than 20% lower above symbionts expressing MaSAMP compared to the GFP control (**Fig. 4B**), the difference was of weak statical significance (χ^2^=2.3; d.f.=1; *P*=0.128).

**Figure 3.**
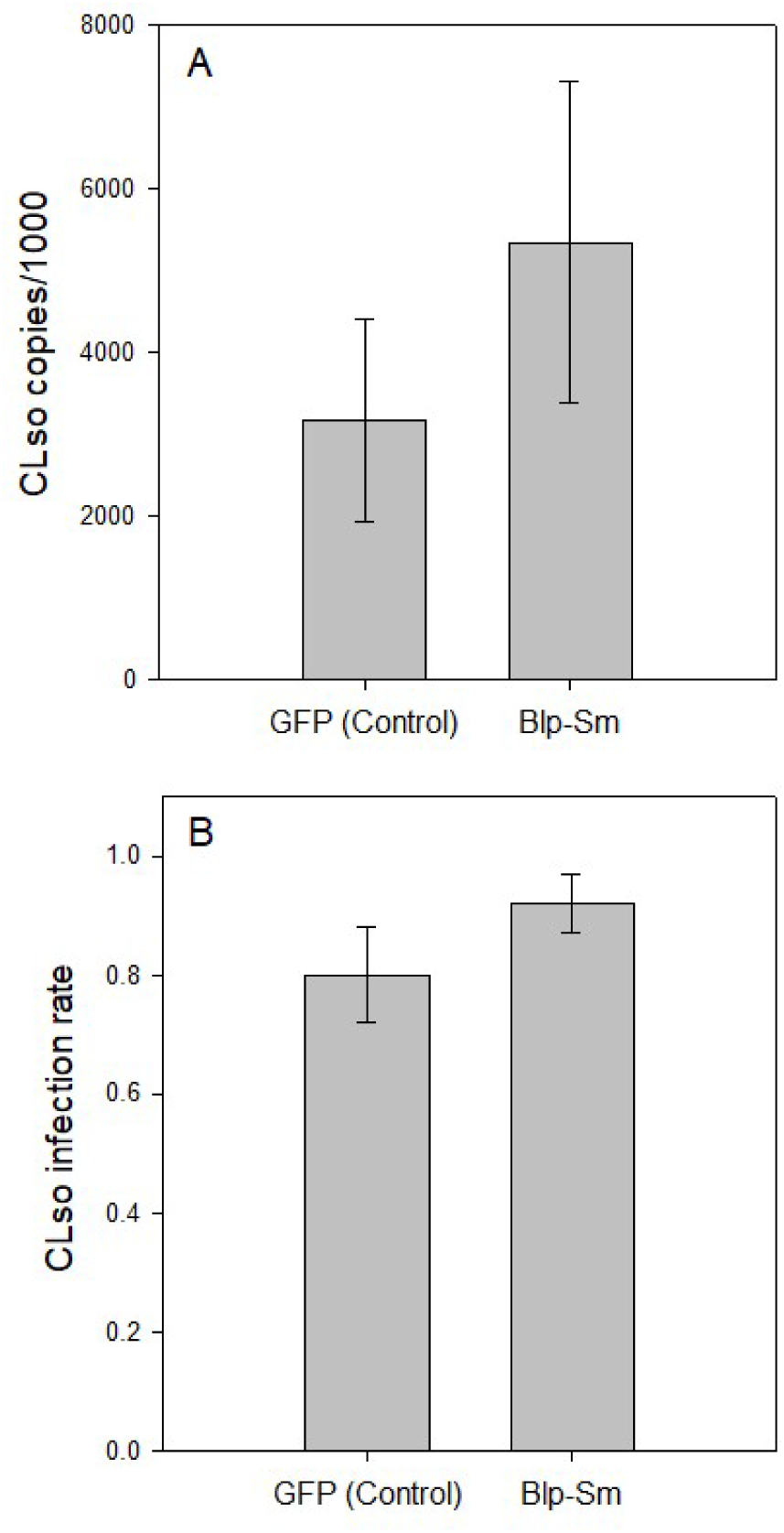
CLso titers (A) and infection rate (B) in stem tissue collected 2-3 cm above symbiont galls producing the antimicrobial peptide, Blp-Sm, versus GFP only (control).

**Figure 4.**
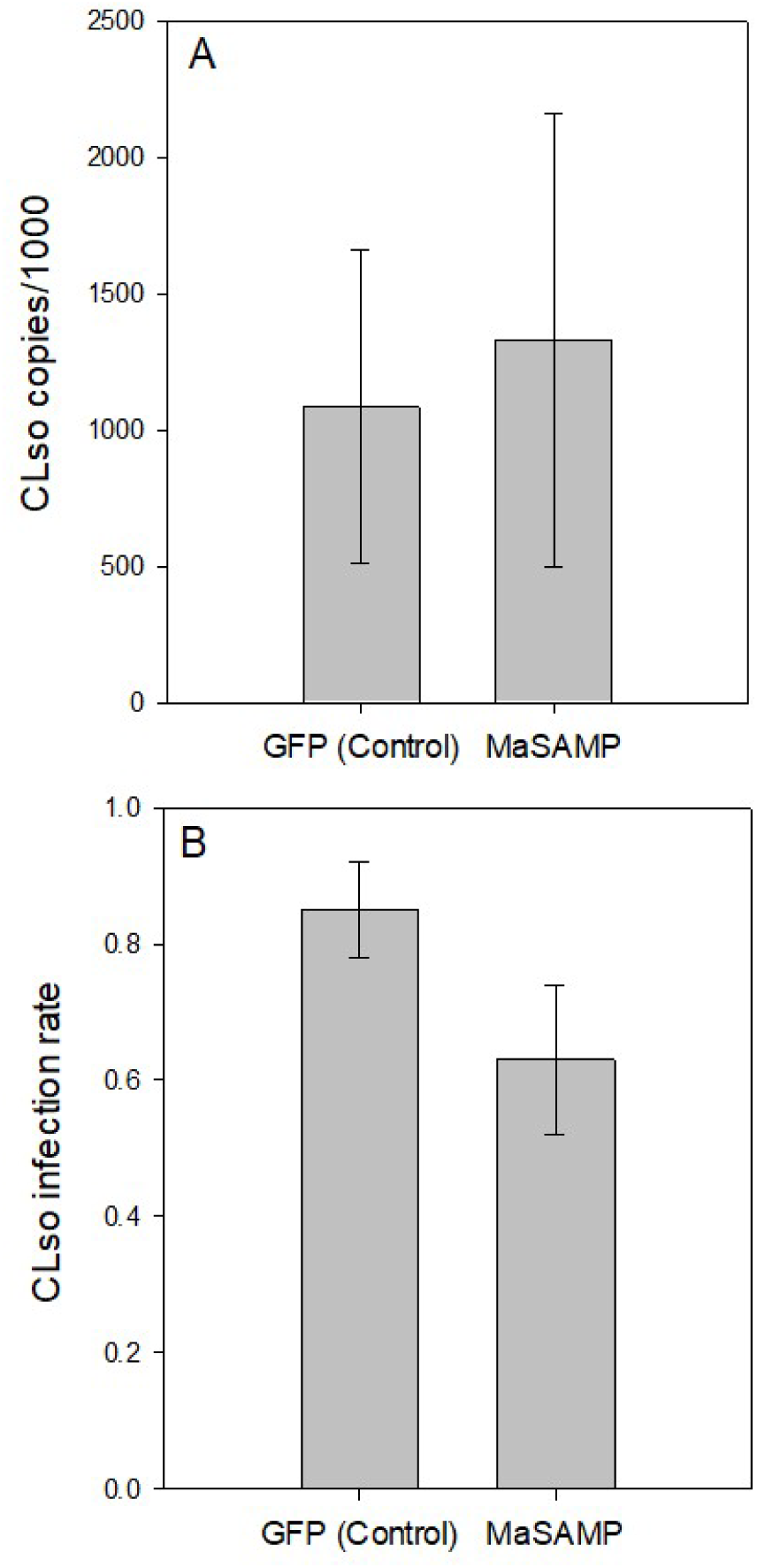
CLso titers (A) and infection rate (B) in stem tissue collected 2-3 cm above symbiont galls producing the antimicrobial peptide, MaSAMP, versus GFP only (control).

**Table 1.**
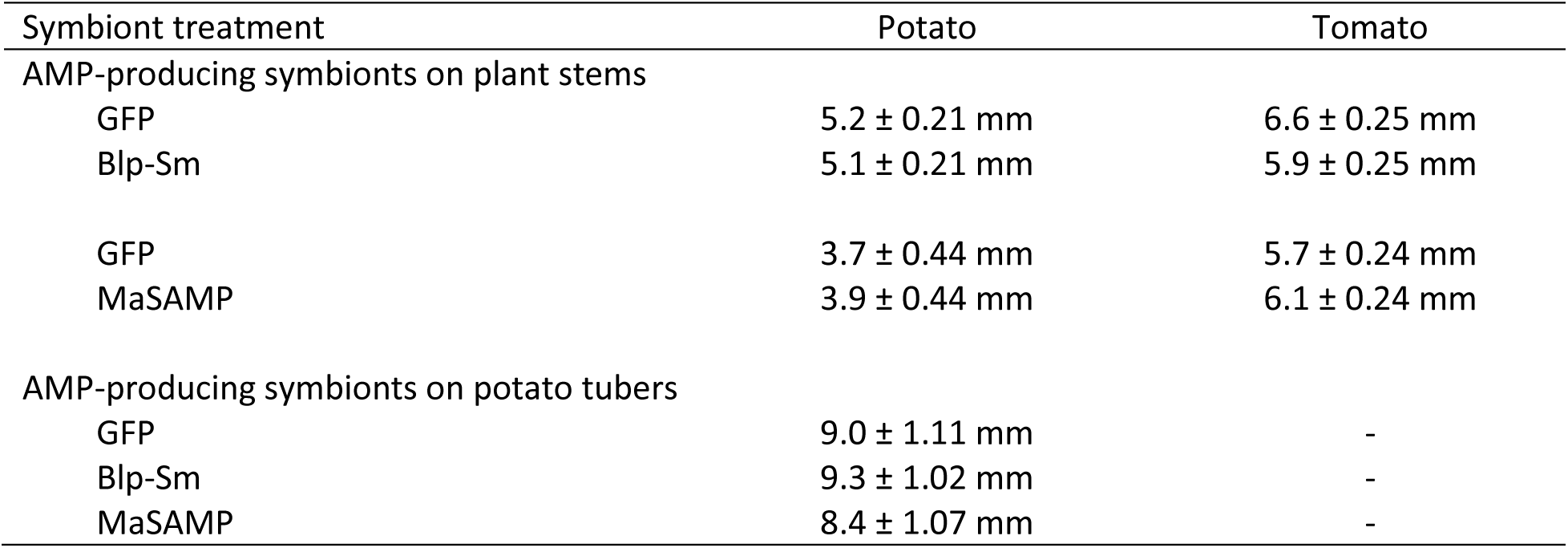
Average symbiont gall sizes (± S. E.) measured at the end of each experiment.

### Symbiont production of antimicrobial peptides reduced CLso accumulation and movement in potato

The potato assay was designed to evaluate whether AMP-producing symbionts affected the movement of CLso between connected potato shoots arising from the same mother tuber (**Fig. 2**). Symbiont galls formed readily on potato stems, typically reaching diameters of 4-5 mm and consistently expressing GFP (**Table 1**).

#### Effects of Blp-Sm-producing symbionts

Blp-Sm-producing symbionts significantly affected CLso titers in both stem and leaf tissues, and these effects differed between source and sink shoots, as indicated by significant treatment x shoot interactions (**Table 2**). In stem tissues, CLso titers in GFP controls were similar between source and sink shoots, whereas titers in sink shoots expressing Blp-SM were nearly undetectable (**Fig. 5A**). In leaf tissues, CLso titers were generally lower in sink shoots than in source shoots, with the lowest titers detected in sink leaves expressing Blp-Sm (**Fig. 5B**).

**Figure 5.**
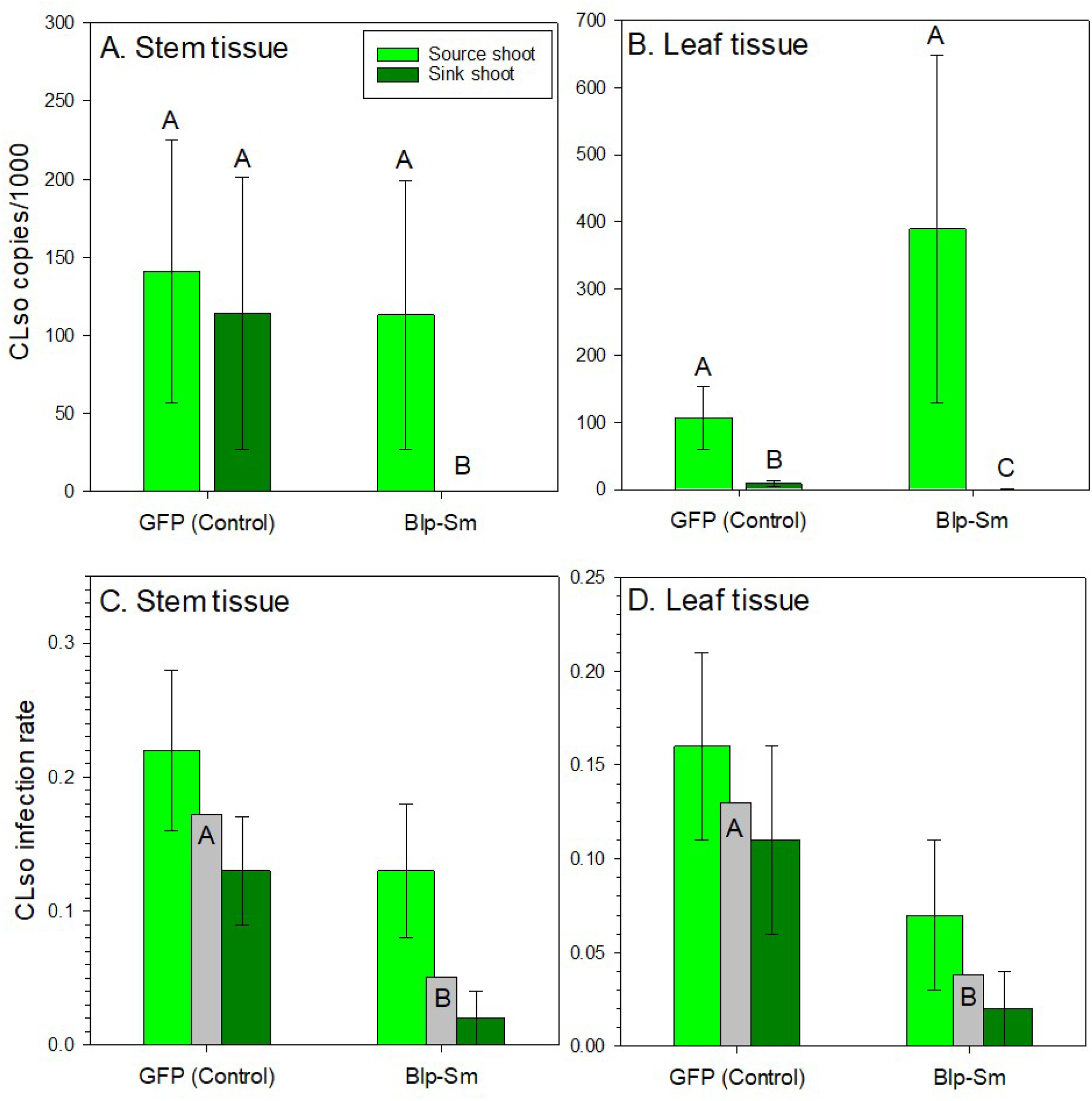
Effects of symbiont expressing Blp-Sm on CLso infection rates in stem (A) and leaf (B) and Clso titers in stem (C) and leaf (D) tissues of source and sink potato shoots. Gray bars represent treatment-level means of CLso infection rates averaged between source and sink shoots. Figure 1 provides a visual description of the experimental design.

**Table 2.** Statistical analyses of CLso titers and infection rates in source and sink shoots of potato plants with symbiont producing the antimicrobial peptides, Blp-Sm and MaSAMP.

| Symbiont treatment | CLso titers |  | CLso infection rates |  |
| --- | --- | --- | --- | --- |
|  | Stem tissue | Leaf tissue | Stem tissue | Leaf tissue |
| **Blp-Sm vs. GFP** |  |  |  |  |
| Treatment | $F=9.3$; $df=1, 19$;<br>$P=0.006$ | $F=2.8$; $df=1, 12$;<br>$P=0.119$ | $\chi^2=3.8$; $df=1$;<br>$P=0.050$ | $\chi^2=4.1$; $df=1$;<br>$P=0.044$ |
| Shoot | $F=9.2$; $df=1, 19$;<br>$P=0.007$ | $F=45.4$; $df=1, 12$;<br>$P<0.001$ | $\chi^2=3.7$; $df=1$;<br>$P=0.050$ | $\chi^2=1.0$; $df=1$;<br>$P=0.310$ |
| Trt x shoot interaction | $F=9.1$; $df=1, 19$;<br>$P=0.010$ | $F=11.6$; $df=1, 12$;<br>$P=0.005$ | $\chi^2=4.6$; $df=3$;<br>$P=0.207$ | $\chi^2=0.4$; $df=3$;<br>$P=0.207$ |
| **MaSAMP vs. GFP** |  |  |  |  |
| Treatment | $F=1.4$; $df=1, 12$;<br>$P=0.256$ | $F=19.5$; $df=1, 10$;<br>$P=0.001$ | $\chi^2=1.2$; $df=1$;<br>$P=0.284$ | $\chi^2=3.2$; $df=1$;<br>$P=0.074$ |
| Shoot | $F=1.7$; $df=1, 12$;<br>$P=0.218$ | $F=25.9$; $df=1, 10$;<br>$P<0.001$ | $\chi^2=0.1$; $df=1$;<br>$P=0.976$ | $\chi^2=0.1$; $df=1$;<br>$P=0.969$ |
| Trt x shoot interaction | $F=0.2$; $df=1, 12$;<br>$P=0.671$ | $F=9.5$; $df=1, 10$;<br>$P=0.012$ | $\chi^2=1.5$; $df=3$;<br>$P=0.673$ | $\chi^2=4.1$; $df=3$;<br>$P=0.251$ |

CLso infection rates were also significantly reduced in plants expressing Blp-Sm (**Table 2**). In stem tissues, infection incidence was lower in Blp-Sm-treated plants compared with GFP controls and was also lower in sink shoots than in source shoots (**Fig. 5C**). In leaf tissues, infection incidence was similarly reduced in plants expressing Blp-Sm compared with GFP controls (**Fig. 5D**).

#### Effects of MaSAMP-producing symbionts

MaSAMP-producing symbionts did not significantly reduce CLso titers in stem tissues, and no treatment x shoot interaction was detected for stem titers (**Table 2**, **Fig. 6A**). In contrast, leaf titers were significantly affected by treatment and differed between source and sink shoots, as indicated by a significant treatment x shoot interaction (**Table 2**). CLso titers in sink leaves expressing MaSAMP were nearly undetectable, whereas titers remained elevated in source leaves and in GFP controls (**Fig. 6B**).

**Figure 6.**
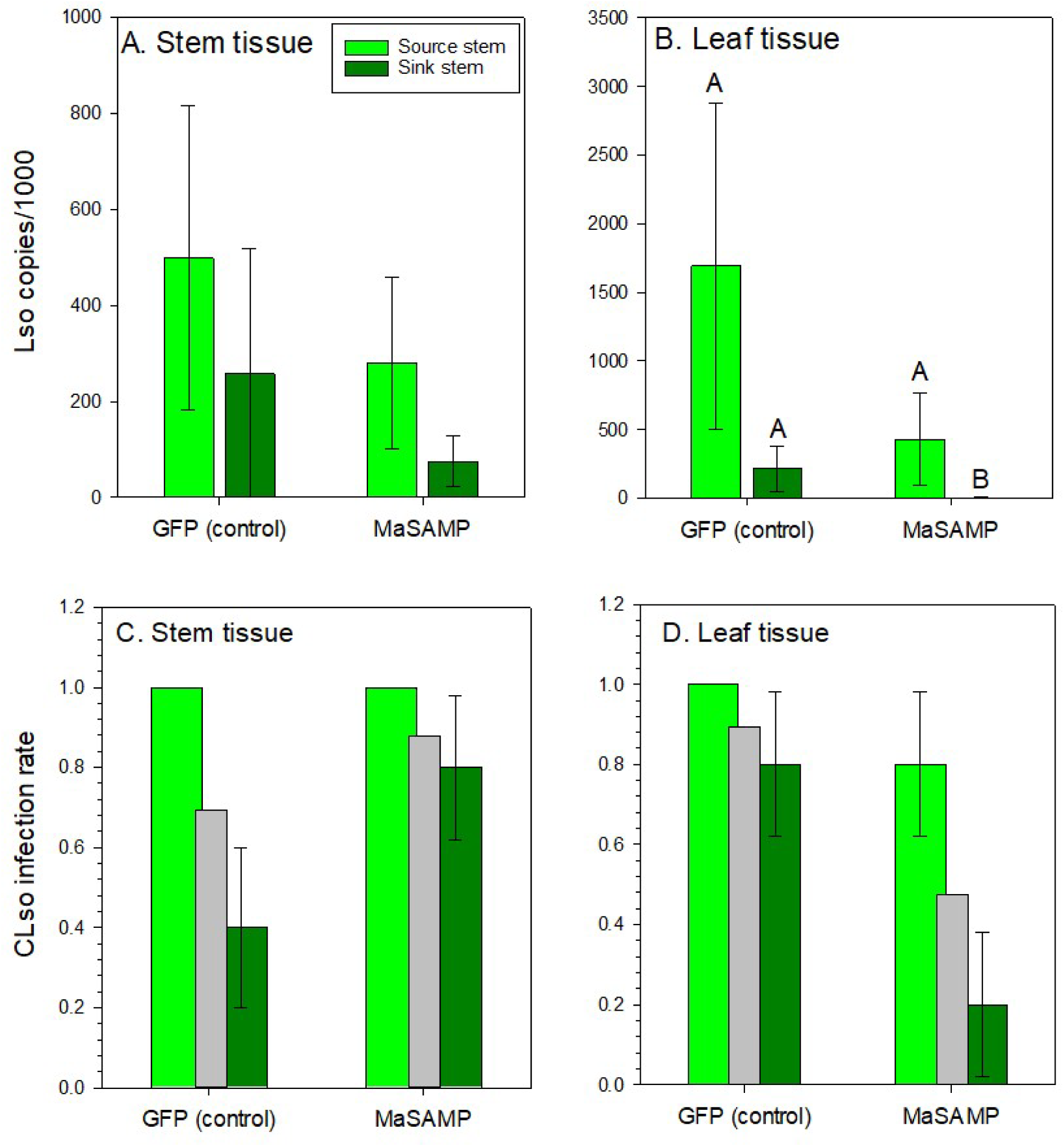
Effects of symbiont expressing MaSAMP on CLso infection rates in stem (A) and leaf (B) and Clso titers in stem (C) and leaf (D) tissues of source and sink potato shoots. Gray bars represent treatment-level means of CLso infection rates averaged between source and sink shoots. Figure 1 provides a visual description of the experimental design.

Infection rates in MaSAMP experiments were generally high across treatments, limiting statistical separation among groups. No significant treatment effects were detected for stem infection rates (**Table 2**, **Fig. 6C**). In leaf tissues, infection rates were numerically lower in plants expressing MaSAMP, particularly in sink shoots, but these differences were not statistically significant (**Table 2**, **Fig. 6D**).

Collectively, the potato assays demonstrated that symbiont-produced AMPs, particularly Blp-Sm, reduced CLso accumulation and infection in sink tissues, consistent with symbiont-mediated inhibition of systemic pathogen movement within plants.

### Seed tuber symbionts expressing MaSAMP reduce CLso titers and zebra chip symptoms

Symbiont galls grew to diameters ranging from 8-9 mm on seed tubers and expressed a high level of GFP (**Table 1**). MaSAMP-expressing symbionts growing on mother tubers of potato plants reduced CLso titers in the plant stems compared with symbionts expressing Blp-Sm or GFP only (**Fig. 7A**; *F*=4.8; d.f.=2, 41; *P*=0.013). In fact, CLso titers in plants with MaSAMP-expressing symbionts were nearly 1/4^th^ of those detected in control plants (**Fig. 7A**). CLso titers in terminal leaves were variable and did not differ among treatments (**Fig. 7B**; *F*=0.48; d.f.=2, 38; *P*=0.625). CLso titers were significantly reduced in tubers near symbionts-expressing MaSAMP compared with the GFP controls and Blp-Sm (**Fig. 7C**; *F*=4.4; d.f.=2, 29; *P*=0.021). Similarly, the proportion of tubers exhibiting zebra chip symptoms was significantly reduced in plants with symbionts expressing MaSAMP compared with GFP controls or Blp-Sm (**Fig. 7D**).

**Figure 7.**
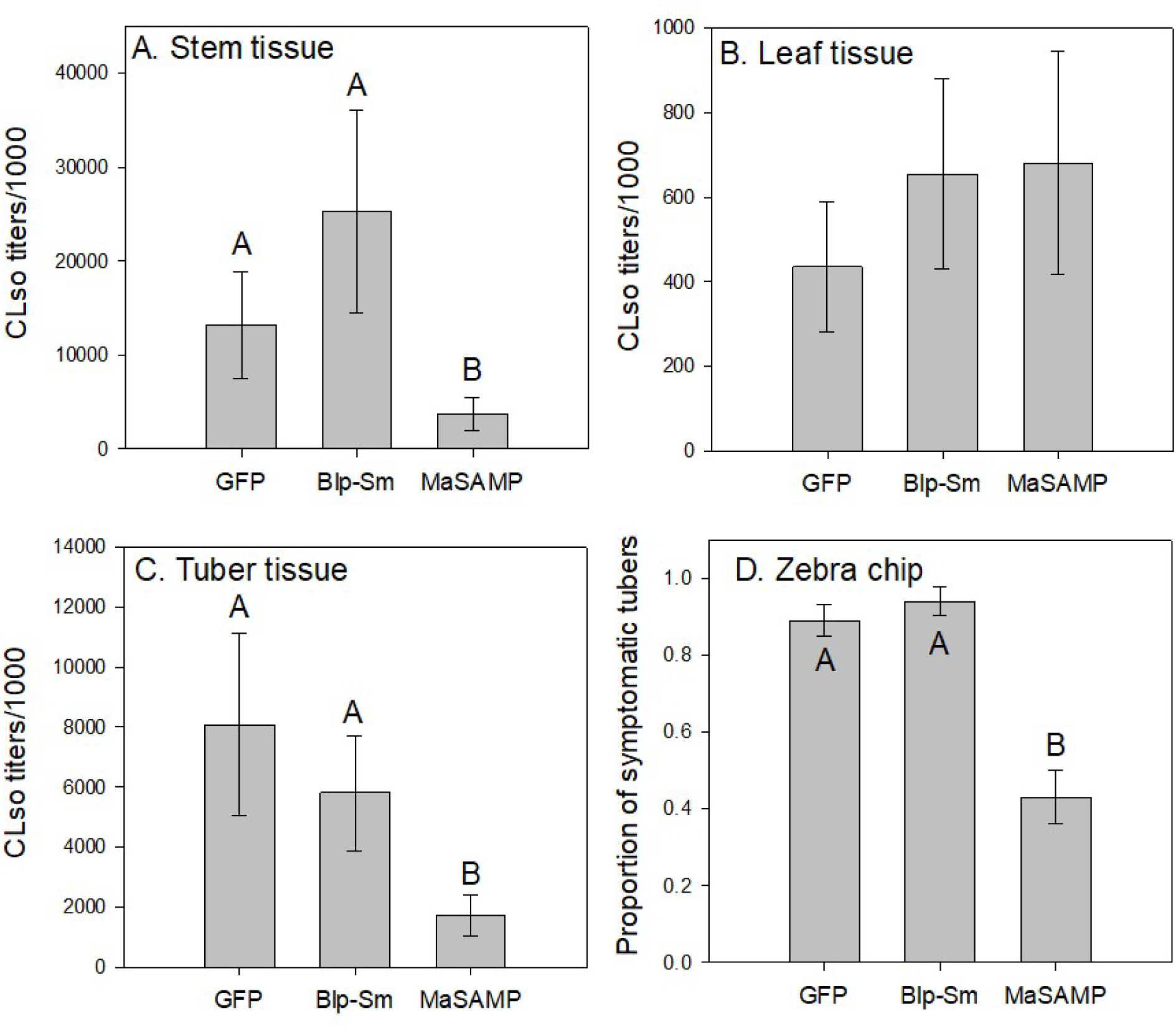
Effects of tuber-inoculated symbiont expressing antimicrobial peptides on CLso titers in stem (A), leaf (B) and tuber (C) tissues, and (D) proportion of symptomatic tubers.

## Discussion

The symbiont concept offers a bioengineering framework for modifying plant phenotypes without whole-plant gene editing (Heck et al. 2026). In this system, a symbiont refers to a reprogrammed plant cell structure that grows externally on the host while remaining structurally integrated with its vascular system. A potential practical application is the production and delivery of antimicrobial peptides (AMPs) to crops infected with fastidious vascular pathogens such as *Candidatus* Liberibacter asiaticus, the presumed causal agent of citrus greening, or *Candidatus* Phytoplasma pruni, which is associated with cherry X-disease. Our results show that AMP-producing symbionts can reduce CLso titers in potato and decrease the incidence of potato zebra chip disease, highlighting the promise of symbionts as a tool for managing vascular pathogens. To test this concept, we engineered symbionts to express either Blp-Sm, a bacteriocin with broad-spectrum antibacterial activity (Dawid et al. 2007, Franz et al. 2007), or MaSAMP, a peptide associated with host resistance to *Ca.* L. asiaticus (Huang et al. 2020). Our experiments evaluated the efficacy of symbiont-produced AMPs on CLso titers both locally in stems near the symbiont gall and distantly in terminal leaves of CLso-infected potato or tomato shoots, as well as in uninfected potato shoots connected to infected shoots through a subterranean seed (mother) tuber.

For AMPs produced by the symbiont to clear infections in terminal leaves on an infected shoot, we hypothesize they must be exported from the symbiont tissue and transported systemically through the plant vasculature. In our experiments, however, we did not obtain compelling evidence for strong or consistent long-distance upward movement of symbiont-produced AMPs into the primary infected foliage. Across both tomato and potato, only slight, non-statistically significant reductions in CLso titers were detected in stems and leaves of shoots directly inoculated with the pathogen. Pathogen titers in these tissues were highly variable, consistent with previous observations of treatment responses in CLso-infected plants (Hunter et al. 2021), yet the underlying causes of this variability remain unclear.

This high variance is characteristic of fastidious vascular pathogens and likely reflects the uneven, patchy distribution of the bacteria within the host phloem architecture (Huang et al. 2020). Further research is needed to identify and control the physiological and environmental factors contributing to variation in CLso titers.

Multiple potato stems arise from a single seed tuber, and because CLso moves readily between these stems via the shared tuber base (Levy et al. 2011), this host provides a useful model system for evaluating whether treatments can inhibit pathogen movement within a plant. Symbiont-produced AMPs may reduce CLso accumulation in opposite sink shoots, i.e., those not directly infected with the pathogen (**Fig. 2**), either by being exported from the gall and transported through the vasculature or by blocking CLso movement from the inoculation site as it passes through symbiont-altered tissues. As expected, infection rates were generally lower in sink shoots than in source shoots regardless of treatment. However, in plants harboring AMP-producing symbionts, CLso was rarely detected in sink stems and leaves. In the relatively few plants where CLso was detected in sink tissues, titers were substantially lower, often approaching zero, compared with sink tissues of control plants. Together, the strongest effects of AMP-producing symbionts were generally observed in sink tissues distal to the primary CLso infection source, consistent with the possibility that symbiont-associated vascular tissues, or the localized accumulation of AMPs near the symbiont base, may function as localized barriers that impede systemic pathogen movement within plants.

Our initial experiments evaluated the effects of AMP production from symbionts growing on plant stems, as described in Heck et al. (2026). In a separate experiment, we inoculated potato seed tubers with symbionts and measured CLso titers in tubers, stems, and leaves. Early attempts to establish symbionts on tuber buds were unsuccessful, likely because tubers were inoculated immediately after removal from cold storage when metabolic activity was low. In contrast, symbiont galls formed readily on tubers that were held at room temperature for at least 24 hours prior to inoculation allowing the buds to break dormancy. These tuber-derived galls were nearly twice the size of those produced on stems and showed relatively high levels of GFP expression. Although expression of Blp-Sm did not significantly affect CLso titers or infection rates, MaSAMP expression led to a significant reduction in CLso titers in stems and tubers and a corresponding decrease in zebra chip symptoms. These results suggest that symptom development in potato tubers may depend more strongly on local pathogen accumulation within tuber-associated vascular tissues than on systemic CLso titers in aerial tissues. These findings further suggest that CLso titers in leaves may not reliably predict pathogen accumulation in tubers or tuber disease severity. Huang et al. (2021) demonstrated that MaSAMP not only exhibits direct antimicrobial activity against Liberibacer species but also induces systemic defense responses in citrus. Thus, the reduced CLso titers and zebra chip symptoms observed in tubers may reflect a combination of direct antimicrobial effects and localized host defense responses induced by MaSAMP-producing symbionts. Critically, our findings suggest that complete suppression of CLso throughout the plant may not be required to reduce economically-important disease symptoms in potato.

Our results are the first to demonstrate the ability to alter plant phenotypes using the symbiont plant engineering concept by delivery of two different AMPs. The two AMPs evaluated in our study did not produce identical patterns of activity *in planta*, suggesting that symbiont efficacy may depend strongly on the biological properties and mechanisms of the expressed peptide. Blp-Sm produced more consistent reductions in CLso movement into sink tissues, whereas MaSAMP was more strongly associated with reduced tuber titers and zebra chip symptoms. These differences may reflect distinct modes of action, including the potential contribution of host defense responses induced by MaSAMP in addition to direct antimicrobial activity. The differing results between the two peptides also suggest that screening assays developed to improve symbiont technology solely focused on titer reduction in leaves may miss effective candidates that successfully suppress target disease symptoms in harvestable organs. Thus, care should be taken to focus future assays on measuring the outcomes most economically meaningful to growers, where feasible. The development and refinement of symbiont technology for managing diseases caused by vascular pathogens, e.g., citrus greening, cherry X-disease, and grape Pierce’s disease, could provide long-term, sustainable control strategies that reduce reliance on pesticides. The ability to inoculate potato seed tubers with symbionts also presents a potential alternative to at-planting neonicotinoid insecticides used to suppress vector-borne vascular pathogens. Future work should focus on increasing the efficacy of AMP production and export from the gall, improving understanding of vascular connectivity between the symbiont gall and host tissues, and enhancing symbiont performance through more potent AMPs or by stacking multiple AMPs within a single construct.

While the symbiont concept remains an early-stage technology, our study further demonstrates that CLso infection in potato provides a rapid and experimentally tractable greenhouse-based model system for optimizing symbiont constructs, evaluating AMP performance and generating insights into vascular pathogen movement that may be adaptable to perennial crops such as citrus and cherry, where disease development and experimental timelines are substantially longer.

## Acknowledgements

The author(s) declared that financial support was received for this work and/or its publication. This research was funded by a grant from the United States Department of Agriculture (USDA) National Institute of Food and Agriculture (NIFA) Project # 2020-70029-33176 and funding from the USDA Agricultural Research Service (ARS) Head Quarters Symbiont Sprint Initiative. MacKenzie Evans, Aspen Scott, Rachel Cook and Doug Harper provided technical assistance.

## Conflict of Interest

MH, MP, and RS are inventors on the patent pending for symbiont biotechnology. There is no symbiont product for sale, and the use of the word symbiont in this paper is for educational and academic purposes only. The symbiont approach is patent pending under U.S. Pat. Appl. No. 17/635, 494: “COMPOSITION AND METHODS FOR MODIFYING A PLANT CHARACTERISTIC WITHOUT MODIFYING THE PLANT GENOME”. The provisional patent application was filed Sept. 20, 2019. The Patent Cooperation Treaty (PCT) was filed Sept. 18, 2020. The US patent application was published Sept. 29, 2022. The USDA does not endorse commercial products, services, or processes by name, trademark, or manufacturer.

The USDA does not endorse linked websites or the information, products, or services they contain.

